# AI supported *in silico* screening of chimeric antigen receptor therapy targets

**DOI:** 10.64898/2026.08.21.746142

**Authors:** Giorgia Moranzoni, Lasse Vedel Jørgensen, Javier Herranz del Cerro, Antoine Andreoletti, Magnus Haraldson Høie, Kristoffer Vitting-Seerup, Mike Bogetofte Barnkob, Lars Rønn Olsen

## Abstract

Chimeric antigen receptor (CAR) cell therapy has achieved transformative clinical success through targeting of CD19 in refractory B cell malignancies, but extension of this strategy to solid tumors, other hematological malignancies, and autoimmune disease has exposed the complexity of target selection. Antigen abundance alone is not sufficient to define a suitable CAR target. Instead, therapeutic efficacy and safety are shaped by a broader set of molecular features, including isoform usage, subcellular localization, secretion, epitope stability, and the structural context in which antibody-derived binding domains engage their target. At the same time, advances in transcriptomics, structural biology, and artificial intelligence (AI)-enabled prediction now make it possible to assess many of these properties systematically. Here, we outline the principal molecular features that characterize effective and safe CAR targets and present a practical framework that integrates public datasets with computational and AI-based tools for their evaluation. Using HER2 as an illustrative case, we show how isoform-resolved expression, single-cell analyses, topology prediction, structure modelling, epitope mapping, and in silico binding analyses can reveal liabilities that are not captured by conventional target-expression screens alone. This framework provides a systematic strategy to prioritize targets and epitopes, guide preclinical investigation, and de-risk clinical translation. We anticipate that such integrative workflows will become increasingly important for moving CAR target discovery from descriptive expression analysis towards informed therapeutic design.

## Introduction

Chimeric antigen receptor (CAR) T cell therapy is a personalized treatment in which patients’ T cells are genetically manipulated to enable cytotoxic killing of cells expressing endogenous surface proteins such as the B-cell receptor CD19. Clinical studies of CD19-specific CARs have shown high efficacy in patients with relapsed and refractory B cell malignancies^1–4^. CAR therapy is now actively explored in many other hematological and solid malignancies (as of time of June 2026, more than 1,000 active CAR cell therapy clinical trials against cancer indications are listed on clinicaltrials.gov) and thus methods for systematic evaluation of novel CAR targets are needed. Recently, CAR T cells have also shown promising results in an autoimmune setting, further indicating the broad setting the therapy is expected to be utilized in^5–7^.

CAR T cells are generated by first harvesting cells by leukapheresis, followed by viral transduction with a CAR construct, thereby modifying the T cells to express a chimeric protein. The CAR itself is a synthetic protein, consisting of an extracellular binding domain, usually an antibody-derived single chain variable fragment (scFv), and the activating signaling domains from the T-cell-receptor complex, including CD3ζ and co-stimulatory domains such as CD28 or 4-1BB. When these CAR T cells are infused back into the patient, they recognize cell-surface proteins through the extracellular domain of the CAR, thereby initiating cytotoxic programs in the T cells, and initiating killing of cells expressing the targeted protein. Because this is a living medicine, it is expected that a majority of treated patients will have residual CAR T cells for the remainder of their lives^8^, and that cells expressing the target protein will therefore be suppressed or removed.

Which cell-surface protein to target is an important component in the design and development of new CAR T cell therapies. Both treatment outcome and persistence of the modified T cells are highly dependent on their ability to sufficiently and continuously engage with the chosen target. Studies have shown that while CAR T cells can lyse cancer cells expressing even low numbers of targets, their ability to proliferate and establish a strong immunological response is dependent on a certain threshold of target density on the cancer cell surface^9,10^. Similarly, downregulation, mutations, and isoform switching of the targeted protein is a major obstacle for successful therapy^11,12^. Finally, expression of the target antigen on non-cancerous tissue must be limited or avoided to alleviate serious on-target, off-tumor toxicities^13^, as these can be life-threatening in a subset of patients.

Recently a number of advanced CARs have been designed to simultaneously target several different cell-surface proteins (called bi-or tri-specific CARs), or rely on multiple input signals for their activation (called logic-gated CARs)^14^. While bi-and tri-specific CARs have been shown to decrease target-escape variants in preclinical models^15–17^, each additional target added to the CAR can result in increased risk of side-effects. A number of groups have also designed CARs that incorporate several input signals to better distinguish malignant and healthy cells. One approach has been to couple targets with sub-optimal signaling domains, such that two inputs are needed for activation^18–20^. Another approach has been to add an inhibitory CAR to the T cells, such that binding to the targeted protein results in inhibition, rather than activation, of the signaling cascade^21^. Because of these advances it has been estimated that there are over 100 different targets amenable for mono-specific CAR therapy and over 100,000 target pairs available for logic-gated CARs^22^. Taken together, there is thus a strong need for workflows that integrate putative targets and target-combinations with suitable data and bioinformatics analyses that assess target properties and druggability.

Several studies have employed bioinformatics analyses for target identification. In one case, publicly available single-cell transcriptomics data from individuals with acute myeloid leukemia (AML) were analyzed, leading to the discovery of CSFR1 and CD86 as potential CAR T-cell therapy targets. These findings were further validated using bulk RNA-seq data using the Genotype-Tissue Expression (GTEx) dataset^23^. Another study explored the AML surfaceome by integrating transcriptomics and proteomics data from malignant and normal tissues. An algorithm was developed to identify targets expressed in leukemia stem cells but not in normal hematopoietic or vital tissues, leading to a promising combinatorial CAR T-cell therapy strategy for AML^24^. In one study, gene expression data from The Cancer Genome Atlas (TCGA) was used to identify up-regulated genes in subtypes of breast cancer. Potential candidates were then evaluated through publicly available proteomics datasets in breast cancer and healthy tissues^25^. Additionally, a targetable landscape for CAR targets was also established by screening clinically tested targets and integrating transcriptional and proteomic data from healthy and cancerous tissues into the analysis^22^. However, the vast majority of studies were based on gene-level target expression, thus ignoring isoform-level expression patterns, which by now is acknowledged to be highly relevant^12,26,27^. As a notable exception, one study by Shaw et al. focused on deploying bioinformatics frameworks to find cancer-specific exons for CAR T-cell therapy targeting^28^, and although annotations such as secretion status were included, we aim here to extend these annotations substantially further.

Recently, artificial intelligence (AI), particularly machine learning and deep learning methods, has emerged as a promising tool in CAR therapy research, aiding in literature mining, predictive modeling, therapeutic efficacy forecasting, and optimization of CAR designs. Current reviews primarily discuss potential benefits and theoretical applications but underscore significant limitations due to dataset quality, model generalizability, and reproducibility challenges^29^. Practical implementations remain scarce, though notable examples include CAR-Toner, an AI-driven tool for optimizing tonic signaling in CAR constructs^30^, and an in silico pipeline integrating AI-guided molecular docking and steered molecular dynamics to accurately select scFv candidates^31^. Here we first provide our perspective of the principal molecular features that characterize optimal CAR targets and then address the technology gap by demonstrating how many existing AI methods can be systematically integrated to computationally evaluate CAR therapy targets currently under clinical investigation.

## Results

### The molecular features of effective and safe CAR therapy targets

In the field of CAR T-cell therapy, target identification has traditionally emphasized expression levels of potential target molecules. However, as clinical applications have expanded, it has become clear that expression alone is insufficient for optimal target selection ^32^. Through a comprehensive review of CAR cell therapy literature, we have identified a number of critical molecular characteristics that define effective and safe CAR targets. These characteristics encompass three fundamental categories: target expression and stability, safety considerations, and binding properties. Given the relative novelty of CAR technology, some of these features lack direct experimental validation in clinical settings. In such cases, we have drawn insights from established antibody-based therapeutic approaches and extrapolated these findings to CAR cell therapy applications. **Figure 1** illustrates these key molecular features and their categorical organization.

**Figure 1.**
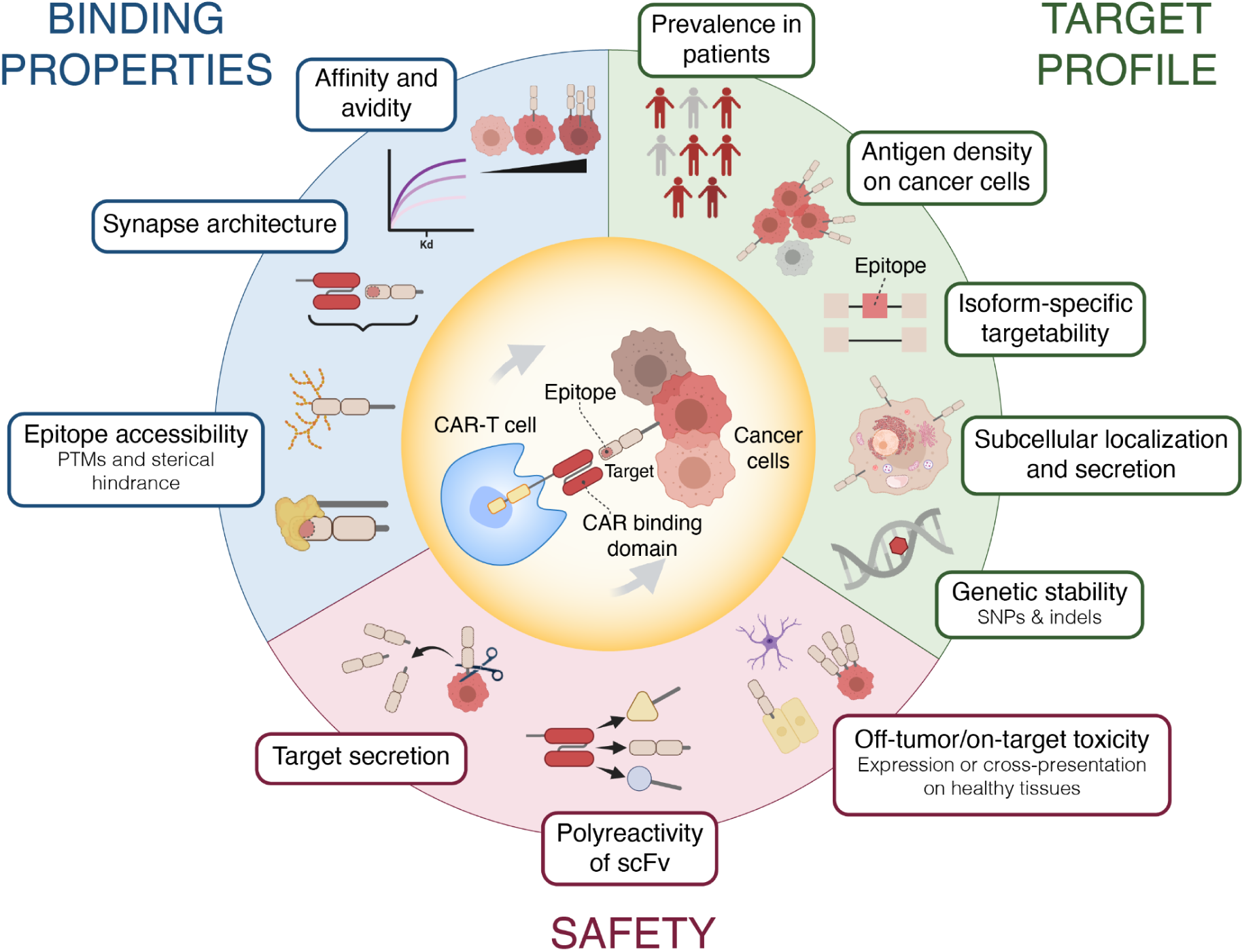
Elements of CAR target selection, subdivided into three major groups: target profile; safety; and binding properties. For target profiling, we consider high target expression in both patient cohorts and antigen expression on cancer cells an important parameter. Target isoforms should be considered individually as they may create decoy targets or be expressed in intracellular locations. Ideally the target epitope should not contain common nonsynonymous SNPs or be subject to isoform switching. Safety is primarily concerned with on-target, off-tumor binding. Likewise, the CAR binding domain should not be poly-reactive and cause off-target binding to other proteins. The target itself should also not be cleavable or able to be cross-presented on other (healthy) cells. Finally, the binding properties of the CAR to the target cell are important: the target and epitope must be available for binding, and not blocked by other protein interactions in cis or trans; or by post-translational modifications (PTMs). CAR size and biophysical properties, such as avidity and affinity, also play an important role in choosing a relevant target, though in silico assessment remains challenging.

### Target expression, isoforms and stability

The primary criterion for selecting targets is the level of expression on both cancer and non-malignant cells. The putative target should be consistently expressed across a heterogeneous population of tumor cells, in the majority of cancer patients, and at levels that allow for recognition by the CAR-T cells. Ideally, targets should be absent on all healthy cells to avoid excessive toxicity^33^. However, targets that satisfy this idealized criterion have yet to be discovered, and all successful CAR therapies thus far have involved some compromise of co-expression on healthy cells.

CAR targets must be cell-membrane proteins, many of which exist in multiple isoforms. Isoforms are different versions of a protein that are produced from the same gene through alternative splicing, and may result in significantly different structures, including in their extracellular domains, and subcellular locations. Alternative splicing leading to isoform switching has been shown to be a common feature in many cancers^34^, and has been characterized as an emerging hallmark of cancer development^35^. Isoform switched proteins might retain essential functions, while the target epitope can be lost. This issue was exemplified in a recent study of the effect on isoform switching on 883 small molecule cancer drugs, targeting 1,434 different proteins. The authors conclude that 76% of these drugs would miss a target isoform if a switch occurred, or induce off-target effects in isoforms expressed in normal tissues^36^.

The efficacy of CAR cell therapy has been observed to be compromised by the “antigen sink” effect, a pharmacokinetic phenomenon where soluble forms of the target antigen sequester the therapeutic cells before they can engage the tumor^37^. While this mechanism is commonly observed with proteolytically shed antigens such as BCMA or Mesothelin, which are released by metalloproteases and γ-secretase^38,39^, alternative splicing represents a potent, genetically driven mechanism for generating these decoys. Isoform switching often leads to the exclusion of transmembrane-encoding exons, resulting in the secretion of soluble variants, such as those observed in PD-L1 resistance models^40,41^. These soluble decoys act as competitive inhibitors that bind the CAR with high affinity, effectively neutralizing the therapeutic dose in the systemic circulation or tumor microenvironment and preventing the formation of the immunological synapse required for tumor elimination^39^.

Conversely, isoform switching can drive resistance by altering the subcellular localization of the target from the cell membrane to the intracellular compartment, thereby rendering the tumor “invisible” to surface-restricted CARs^42^. A definitive example of this mechanism is observed in B-cell acute lymphoblastic leukemia (B-ALL) with the CD19Δex2 isoform. Research indicates that the splicing-mediated exclusion of exon 2 results in a misfolded protein that fails to associate with necessary transport chaperones like CD81, leading to its retention in the endoplasmic reticulum (ER) rather than trafficking to the cell surface^43^. This shift creates a phenotype where the tumor remains transcriptionally positive for the antigen but epitope-negative, allowing malignant cells to escape CAR T cell recognition and drive relapse despite the continued intracellular expression of the target^44^. For these reasons, we have previously posited that CAR target considerations should start at the isoform level^12^.

Lastly, the epitope on the targeted protein should be genetically and structurally stable. Genetic instability can be caused by nonsynonymous single-nucleotide polymorphisms (SNPs) or variants either directly impacting the epitope sequence or the availability of the epitope by imposing structural changes, thus affecting the binding affinity of the scFv domain of the CAR, as has been seen with antibody-based therapies^45^.

### Safety

Safety is another major consideration when exploring CAR targets. What constitutes a good target is highly dependent on which cell-type is targeted and how dependent tissue and organ homeostasis is on the given cell-type. Safety considerations are often closely tied to target expression and since most proteins are also expressed in healthy tissues, this can increase the risk of so-called on-target, off-tumor toxicities. Still, the CAR field has shown that biomedical engineering can overcome many of these challenges, once they are known.

As an example, we can consider two CAR targets, CD19 and CD7, which are primarily expressed on B-and T-cells, respectively. CD19 is the most successful CAR target in clinical use today^46^, even though it is broadly expressed across multiple healthy B-cell subsets, in addition to their malignant deviates. Still, targeting the majority of B cells is acceptable, since B cell loss can be mitigated by immunoglobulin substitution^47^. Conversely, while CD7 represents an intriguing target in T-cell acute lymphoblastic leukemia (T-ALL)^48^, CD7-directed CARs have been shown to cause immediate deletion of both endogenous T cells as well as CAR T-cells themselves (so-called fratricide), because the target is broadly expressed on healthy T cells^49,50^. Besides inhibiting the persistence of the CAR T cells, this T cell aplasia cannot easily be substituted, thus resulting in the serious risk that patients will be susceptible to life-threatening infections following therapy^51^. For these reasons, a major emphasis has been put on identifying target antigens that are tumor-specific.

However, this is often easier said than done, due to the ability of CAR T cells to react to and kill cells expressing even a few copies of the target proteins. Many antigens that were thought to be tumor-specific have thus turned out to be expressed at low levels on normal tissue as well, and therefore lead to off-tumor effects. These on-target/off-tumor effects have in some cases resulted in serious toxicities and fatalities^52^. For example, CD19 has also been detected on a rare population of pericytes in the brain microcirculation, which has led to the speculation that this might play a role in the neurotoxicities associated with CD19-specific CAR treatments^53^. A more prominent example of on-target/off-tumor effects is the breast cancer marker, HER2, which is highly expressed in certain breast cancer subtypes, but also by cells in healthy tissues, including the heart, lungs, and the central nervous system. Targeting it using CAR T-cells caused fatal off-tumor effects in one trial^54^. Similarly, CAR T-cells directed towards mesothelin, carbonic anhydrase IX (CAIX) and CEACAM5 have also been shown to cause on-target/off-tumor toxicities^55–57^. In the case of mesothelin, the protein was only upregulated in benign lung epithelial cells following lung-injury^55^, further highlighting the challenges associated with identifying expression in healthy tissue. As is clear from the above examples and discussion, a detailed and broad understanding of which cell-types express a given antigen is thus a key priority when assessing the safety of a target.

Non-specific binding of CAR paratopes to off-antigen epitopes theoretically represents another safety challenge^58^. As has been seen with antibody-based therapeutics^59^, such interactions may occur when the paratope has low specificity, leading to unintended activation of CAR T cells by structurally similar but unrelated proteins^60,61^. While not seen in clinical trials, this off-target activity could theoretically trigger toxicities, including damage to healthy tissues, and reduce therapeutic efficiency by diverting effector functions away from tumor cells, as has been seen with other T-cell therapies^62,63^. To minimize these risks, paratope engineering strategies aim to enhance binding specificity while maintaining high affinity for the intended target. Approaches such as optimizing binding domains through mutagenesis ^64^, employing affinity maturation techniques^65^, and integrating dual-antigen recognition systems have shown promise in mitigating off-target binding.

Finally, secretion or membrane cleavage of the targeted protein should also be considered as a safety concern, as secreted proteins could lead to non-specific, off-target response if the protein is bound or endocytosed by healthy cells. This concern has also been raised for antibody-drug conjugates^66–68^. Additionally, secreted proteins have been shown to inhibit CAR T cells from functioning in some instances, by blocking the binding domain of the CAR before it reaches the target cell. For example, CARs targeting Igκ light chain, have reduced killing efficacy due to the secreted nature and high abundance of this target in plasma^69,70^. Similar effects have been seen with CARs targeting VEGFR-2^71^. In other cases, such as CARs targeting CD30 and mesothelin, soluble variants of the proteins did not inhibit CAR T-cell function^72,73^. Still, CARs can be made to react towards soluble ligands if the CAR itself can dimerize in the cell-membrane^74^, and so whether a target which is also soluble can be accepted, will also be dependent on the CARs structure and binding properties, which we will review next.

### Binding properties

While target expression in malignant and benign tissues remains a key criterion for CAR T-cell target selection, recent investigations have underscored additional molecular nuances in CAR:target interactions, which we collectively referred to here as binding properties. For example, it has become clear that how CAR T cells bind to tumor cells and form synapses play an important role in both immediate CAR activation and persistence of T cells^75,76^.

PTMs are chemical alterations that proteins undergo after synthesis, influencing their structure and function. In the context of CAR T-cell therapy, glycosylation - the addition of sugar moieties to proteins - can modify the target antigen’s structure, potentially hindering CAR binding. Heard et al.^77^ provided a structural analysis of how glycosylation affects the interaction between CAR T cells and CD19. They showed that specific glycosylation patterns on CD19 can alter the epitope’s conformation, thereby affecting the binding affinity and specificity of CAR T cells. PTMs can also create steric hindrance that obstructs the accessibility of the CAR paratope to the target epitope. For example, the tumor-associated mucin MUC1, which forms dense, glycosylated structures on the cancer cell surface. These extensive glycoprotein networks can thus create a physical barrier, obstructing CAR access to specific epitopes. A CAR targeting MUC1 was engineered to overcome these challenges, by incorporating a flexible IgD hinge region to enhance the receptor’s reach and flexibility, thereby improving its ability to navigate the steric hindrance presented by the MUC1 glycoprotein layer^78^.

Traditional CAR T cells (utilizing either CD28-CD3z or 4-1BB-CD3z) have been shown to require 10-100 fold higher antigen-density to become activated, than via T cell receptor (TCR) binding to peptide-MHC (pMHC)^79,80^. Additionally, many CAR:target interactions were found to be unproductive or result in sub-lethal interactions^81^, underscoring key differences from TCR-initiated cytotoxicity. These observations can partly be explained by poorer signaling capacity of the CAR construct^82^ and a decreased ability to bind other co-accessory proteins on target cells^83^. However, it is also clear that CAR T cells do not always form a canonical immunological synapse^84,85^, possibly due to differences in how mechanical forces and microclusters arise between the CAR and target molecules^86,87^, as seen with traditional TCR:pMHC interactions^88^. Taken together, the affinity (K_D_) between CAR and target protein, as well as the overall avidity between CAR and target cells, thus likely play a key role in CAR cytotoxicity. Indeed, in a review of traditional CARs targeting solid tumor targets, CARs with an CAR:target K_D_ >100 nM, demonstrated no clinical responses^89^. Conversely, affinity-tuned CAR T cells might possibly be used to avert off-target binding in healthy tissue with low expression of a target, as was recently done with HER2-specific and mesothelin-specific CAR T cells^90,91^ and with CD5-specific CARs, to avoid fratricide^92^. As such, understanding the number of target molecules per cell of a putative target can become quite important.

Finally, structural features of the CAR itself influence therapeutic efficacy. Studies have shown that adjusting the spacer length between the scFv and the transmembrane domain can significantly impact CAR functionality, likely by optimizing contact with epitopes positioned at varying distances from the cell membrane^93–99^. As such, a clear understanding of both target protein structure and whether the epitope is distally or proximal to the membrane are important.

Collectively, these insights emphasize that, in addition to target expression, a comprehensive understanding of antigen modifications, spatial presentation, and CAR architecture is paramount for maximizing tumor cell recognition and activation.

### Systematic *in silico* evaluation of targets

Several large-scale screens and curated target landscapes have already proposed numerous candidate antigens for CAR cell therapy across hematological and solid malignancies^22,23,24,25^. Rather than aiming to nominate additional targets, we here present a framework, consisting of publicly available databases and computational tools, to systematically evaluate the suitability and liabilities of putative antigen targets for CAR T-cell therapy. While we attempt to take most features reviewed above into account, not all of them can be currently assessed computationally. Here, we exemplify the analysis by assessing the targetability of HER2, as this antigen has been explored in multiple different CAR trials (as of June 2026, 51 completed or ongoing trials with HER2 CAR cells against a cancer indication are listed in clinicaltrials.gov). While this workflow is designed to perform deep evaluation of individual targets, the analyses can in principle be parallelized to perform broader screens. **Figure 2** visualizes an analysis workflow with the proposed steps of evaluation. Vignettes outlining code for major analysis pipelines, as well as a GUI walk-through of selected analyses that can be performed without coding are available at: https://biosurf.org/cart.html.

**Figure 2.**
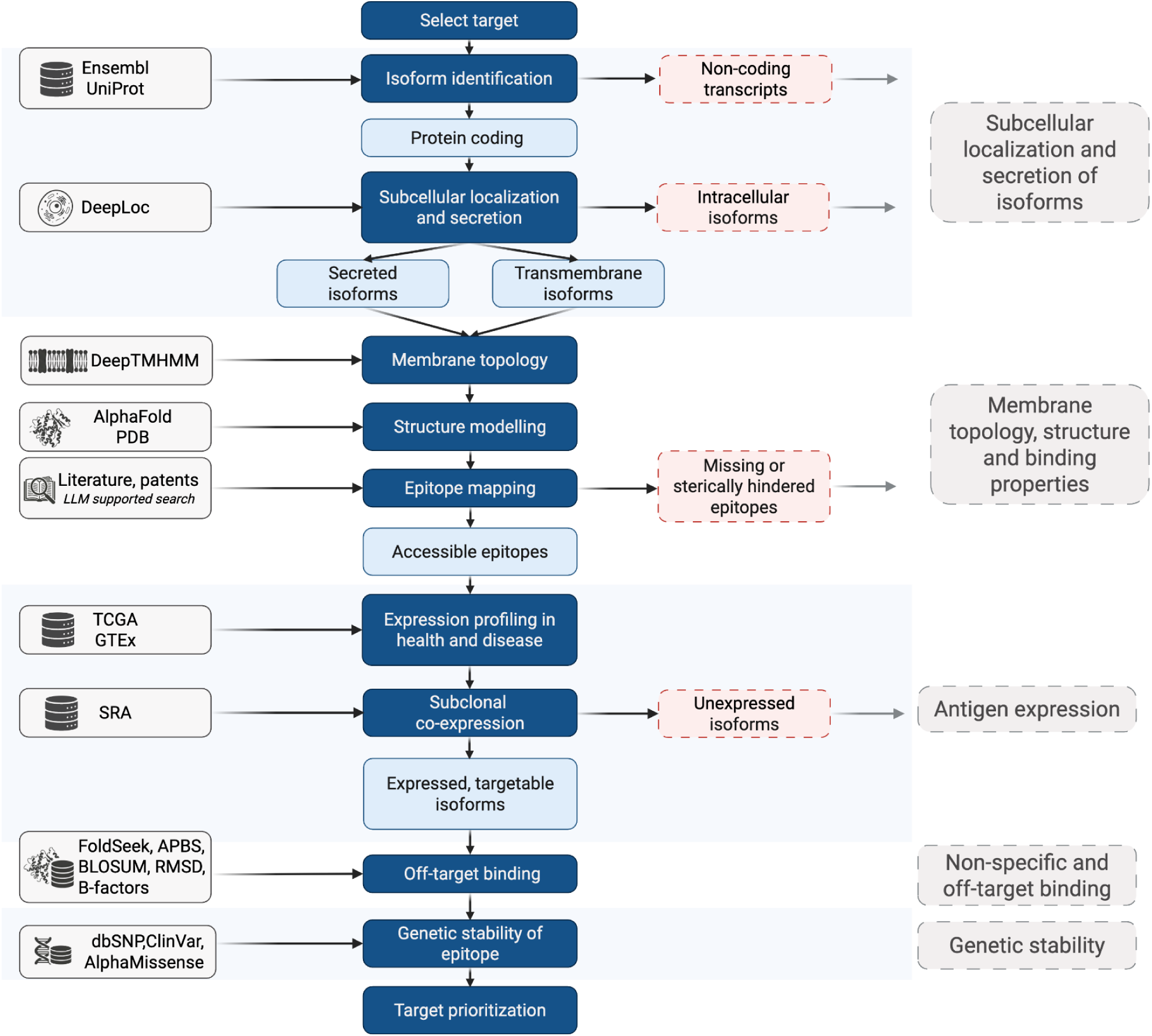
Workflow for systematic in silico evaluation of candidate CAR targets. Protein-coding isoforms are first identified from Ensembl or UniProt and classified by predicted subcellular localization and secretion using DeepLoc. Transmembrane isoforms are then evaluated for membrane topology, three-dimensional structure, and epitope retention using DeepTMHMM, AlphaFold, and literature-or patent-derived epitope information, enabling distinction between accessible and sterically hindered or buried epitopes. In parallel, transcript expression is assessed across healthy and malignant tissues using TCGA and GTEx, and subclonal co-expression is evaluated in single-cell datasets from the Sequence Read Archive (SRA), to identify expressed, targetable isoforms. If epitopes are known, FoldSeek and AntiFold can be used to evaluate potential off-target binding to structurally similar proteins, while dbSNP, ClinVar, AlphaMissense, and AntiFold can be used to assess the effects of genetic variation on epitope stability and recognition. The combined analyses support evidence-based prioritization of targets and epitopes for preclinical CAR development.

### Subcellular localization and secretion of isoforms

For antigens to be targetable using CAR cell therapy, they must localize to the cell membrane. The subcellular location of canonical proteins, isoforms, as well as common and somatic variants can be predicted using algorithms, such as DeepLoc2^100^. The first step in our analysis was to extract all isoform amino acid sequences, predict their subcellular localization and divide them into three broad categories: cell membrane-bound isoforms, secreted isoforms, and intracellular isoforms. The cell membrane-bound isoforms may serve as CAR targets, while isoforms exclusively localizing to intracellular compartments should be excluded from further analysis. Secretion can reduce the availability of cell surface antigens, potentially resulting in antigen escape^38^. Moreover, the secreted isoforms may remain in the extracellular space, binding to and blocking the CAR, thereby serving as decoys hindering its interaction with any membrane-bound isoforms^39^. Consequently, epitopes present in secreted isoforms might be disadvantageous as a targeting site, prompting a search for epitopes unique to cell membrane-anchored isoforms. As shown in **Figure 3A**, 8 isoforms are predominantly localized to the cell membrane, 4 are predominantly secreted, while none of the isoforms of HER2 exclusively localize in intracellular compartments. It should be noted that an isoform can localize to multiple different subcellular compartments, and that DeepLoc2 provides probabilistic predictions for all compartments.

**Figure 3.**
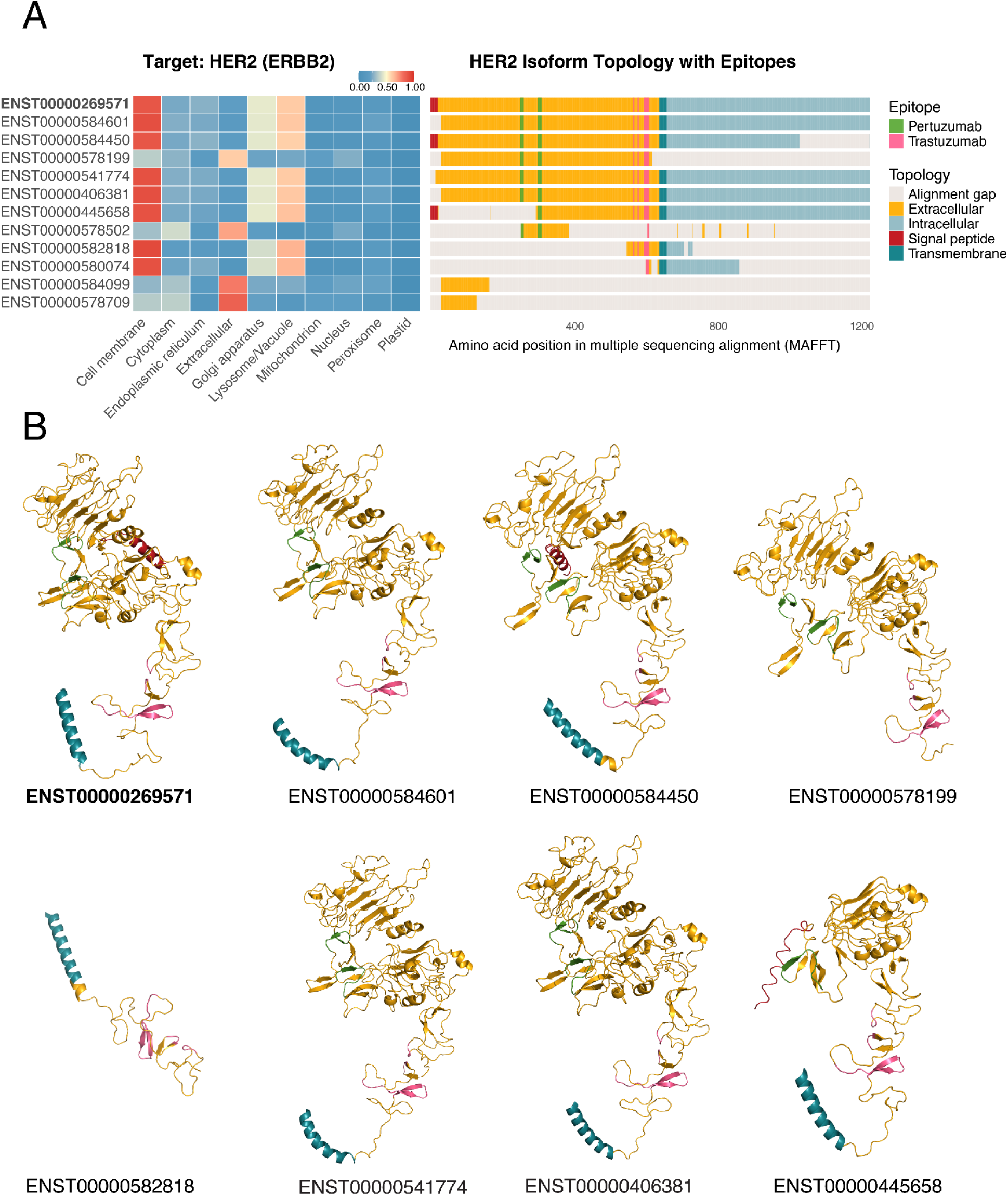
A) The subcellular location predictions of different HER2 (*ERBB2*) isoforms using the DeepLoc2 algorithm (left) as well as topology predictions of each of the isoforms using the DeepTMHMM algorithm (right). The epitope sequences of both trastuzumab and pertuzumab are highlighted in pink and green, respectively. **B)** AlphaFold3-predicted protein structures of the eight HER2 (*ERBB2*) isoforms that contain at least one of the trastuzumab (pink) or pertuzumab (green) epitopes. The endodomains are omitted for clarity. ENST00000578709, ENST00000584099, ENST00000582818, and ENST00000580074 were omitted as they do not contain the full epitope sequence for neither pertuzumab nor trastuzumab.

### Membrane topology, structure, and binding properties

The topology and ectodomain structure of membrane-anchored isoforms are important features of potential targets to evaluate targetability and potential cross-reactivity. We predicted the topology of each expressed HER2 isoform using DeepTMHMM^101^ (**Figure 3A**). Eight different isoforms are predicted by DeepTMHMM to contain a transmembrane domain, while 4 other isoforms lack the transmembrane domain and endodomain. Two isoforms (ENST0000084099 and ENST00000578709, in the bottom rows of **Figure 3A**) appear heavily truncated and were excluded from structural analysis. On all isoforms, we then mapped the epitopes for trastuzumab^102,103^ and pertzumab^104^ - the two primary anti-HER2 treatment options. Both antibodies have been approved by the U.S. Food and Drug Administration (FDA) for the treatment of HER2-positive breast cancer and other HER2-driven cancers^105^. Trastuzumab binds to extracellular domain IV of HER2 and works by inhibiting downstream signaling and mediating antibody-dependent cellular cytotoxicity (ADCC). Pertuzumab binds to extracellular domain II of HER2 and works by preventing receptor dimerization (particularly HER2/HER3) and thereby blocking ligand-induced signaling^105^.

We found the trastuzumab epitope to be present in 7 out of 8 membrane-bound isoforms (the last membrane-bound isoform has a significantly truncated ectodomain), and in 1 out of 4 secreted isoforms. The pertuzumab epitope was found to be present in 5 membrane-bound isoforms and 1 secreted isoform (note that ENST00000578502 may appear to contain the pertuzumab in **Figure 1A**, but the full epitope sequence is, in fact, truncated). Next we predicted the 3D protein structures of all 8 isoforms containing at least one of the epitopes using AlphaFold3^106^, to compare the structural epitope availability. In all isoforms where the full epitope sequences were present, we found no indications that binding would be sterically differentiated across isoforms (**Figure 3B**).

### Antigen expression

Next, we examined the expression of all membrane-anchored and secreted transcripts in bulk RNA sequencing data. This data type provides a broad overview of transcript expression and is available for a large number of patient tumor samples and healthy donor tissues. HER2 is often overexpressed in solid tumors and is an established target for monoclonal antibodies in breast and gastric cancers^107^. A number of HER2-specific CARs are being tested in clinical trials, including against glioblastoma multiforme (GBM)^108,109,110,111^, and breast cancer^112^, and tested preclinically in additional indications, such as stomach adenocarcinoma^113–115^. The expression of HER2 in healthy tissues should be considered carefully, as off-tumor targeting of HER2 by CAR T cells can lead to severe adverse effects. Notably, a fatal case was reported where a patient died due to CAR T-cell therapy attacking HER2-expressing healthy lung tissue^54^.

Transcript-level expression patterns for the eight epitope harboring isoforms are illustrated in **Figure 4A**, and provides a comparative analysis of HER2 isoforms across healthy tissues from GTEx project (for visualization purposes pooled in three risk categories: “critical”, “important”, and “other tissues”) and three cancers from the Cancer Genome Atlas (TCGA): GBM (n = 166), breast invasive carcinoma (n = 1099), and stomach adenocarcinoma (n = 414). The two full-length membrane-bound isoforms, ENST00000269571 and ENST00000541774, exhibited the highest abundance across all tissue categories. While it is elevated in breast invasive carcinoma and stomach adenocarcinoma, its substantial expression in healthy vital and important organs underscores the inherent risk of on-target, off-tumor toxicity previously observed in clinical settings.

**Figure 4.**
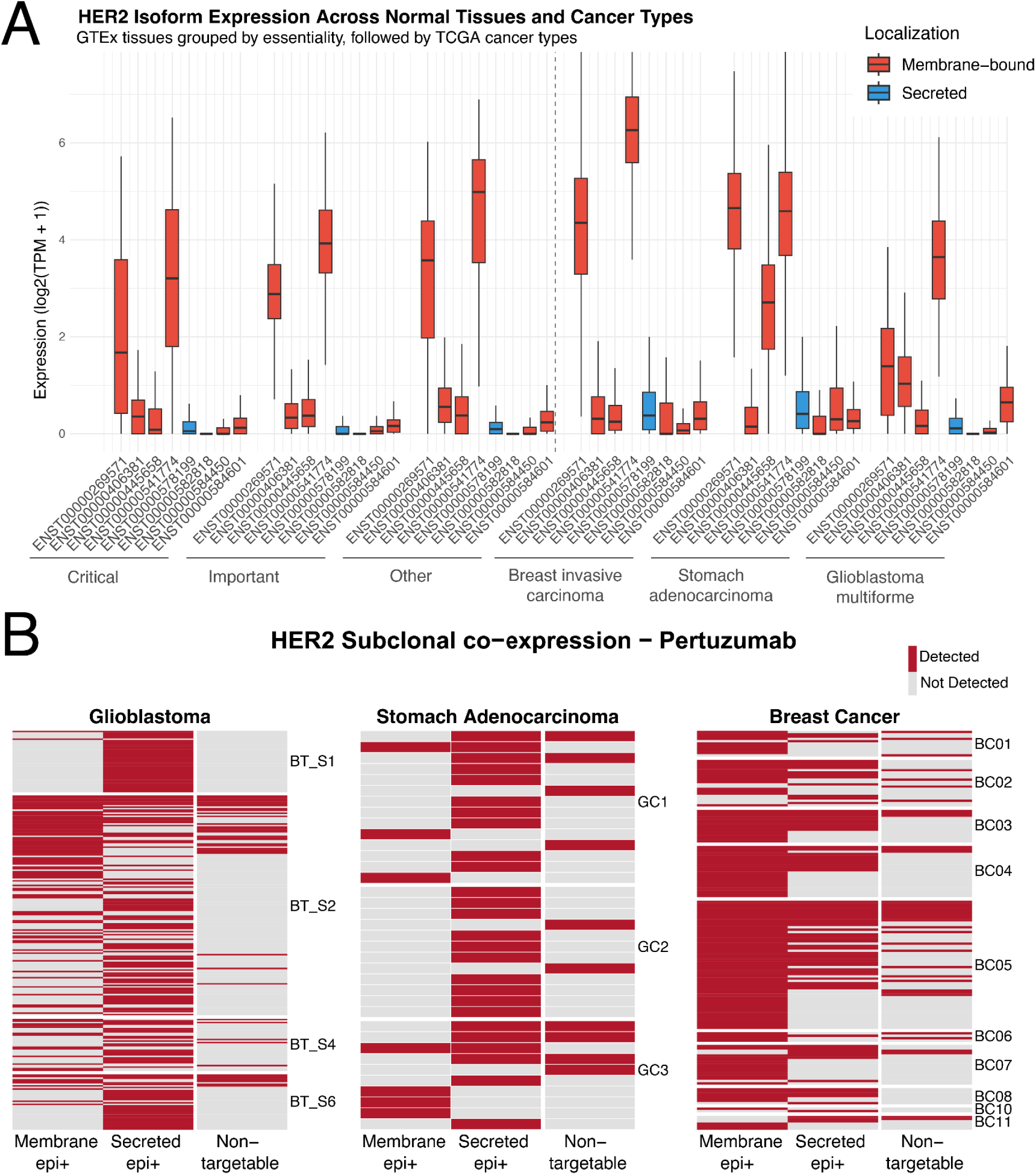
A) The RNA expression of the eight epitope-harboring transcripts. The expression of each transcript was extracted from TCGA and GTEx datasets obtained from the UCSC Xena Browser. Expression in healthy tissues were pooled into three broad categories for legibility (critical: brain, blood, blood vessel, heart, liver, lung, and nerve; important: pancreas, kidney, stomach, colon, small intestine, and bladder; other: spleen, uterus, ovary, testis, prostate, breast, and skin). **B)** RNA co-expression binary heatmap of HER2 isoforms across single cells with detectable HER2 transcripts in glioblastoma, stomach adenocarcinoma, and breast cancer. Each row represents an individual cell, grouped by patients; columns indicate detection (red) or absence (grey) of at least one transcript classified as membrane-bound epitope-positive (5 transcripts), secreted epitope-positive (2 transcripts), or non-targetable (5 transcripts) for pertuzumab. Abbreviations: TPM, transcripts per million. TCGA, The Cancer Genome Atlas Program. GTEx, Genotype-Tissue Expression.

Notably, the truncated membrane-bound isoform (ENST00000445658) shows a more restricted expression profile; it remains expressed at low levels across most healthy tissues, but is expressed at higher levels in the stomach adenocarcinoma samples, than in healthy tissue samples. The trastuzumab epitope is conserved across all expressed membrane bound isoforms, while the pertuzumab epitope is not fully present in this isoform, and would miss it altogether. While neither antibodies can be used to selectively target the overexpressed isoform, its presence suggests that isoform-level screening can be used to identify potential therapeutic windows where a specific cancer-associated variant may be targeted to improve safety. Finally, the secreted isoforms were generally detected at low transcript levels across both normal and cancerous tissues, suggesting that their contribution as soluble decoys or antigen sinks may be limited, although this cannot be determined from transcript abundance alone.

Bulk RNA sequencing provides an overview of isoform expression across cohorts, but cannot determine whether targetable and untargetable isoforms are co-expressed within the same malignant cells or distributed across distinct subclones. This distinction is important, as even a small population of tumor cells lacking the targeted epitope may evade CAR T-cell killing and expand under therapeutic pressure. To assess isoform expression at single-cell resolution, we analyzed Smart-seq2 data from GBM (n = 4 patients, 913 cells), breast cancer (n = 10 patients, 171 cells), and stomach adenocarcinoma (n = 3 patients, 60 cells), restricting the analysis to malignant cells. For each cell, reads were pooled into three categories: membrane-bound isoforms harboring an epitope (5 transcripts), secreted isoforms harboring an epitope (2 transcripts), and untargetable isoforms lacking the epitope (5 transcripts).

Across the three cancers, a fraction of tumor cells showed no detectable HER2 transcripts at all (GBM: 72.29%, breast cancer: 7.60%, stomach adenocarcinoma: 40%). These cells may truly lack HER2 expression, expressing HER2 below the detection limit of single-cell RNA sequencing. The three cancers had varying proportions of membrane-bound, epitope positive cells with detectable HER2 transcripts (GBM: 39.92%, breast cancer: 89.87%, stomach adenocarcinoma: 19.44%), indicating that breast cancer would in principle be the most suitable cancer for HER2 targeted therapy (**Figure 4B**). For GBM and adenocarcinoma, the majority of cells expressed only secreted, epitope positive isoforms, which is not ideal for targeting (GBM: 57.71%, breast cancer: 8.23%, stomach adenocarcinoma: 66.67%). A small fraction of cells expressed exclusively untargetable isoforms lacking the epitope (GBM: 2.37%, breast cancer: 1.90%, stomach adenocarcinoma: 13.89%). While rare at baseline, such cells may still be clinically relevant, as selective pressure from therapy could favor expansion of untargetable subclones, analogous to isoform-mediated escape observed for CD19-targeted CAR T-cell therapies. The same analysis was performed for the trastuzumab epitope, yielding similar overall results (**Supplementary Figure 1**).

### Non-specific binding and off-target effects

Another aspect to consider is the identification of non-specific binding, which may contribute to off-tumor toxicities. This effect may occur if the scFv fragment of a CAR binds epitopes on structurally similar proteins. In principle, this could occur for any target protein and may cause toxicity if the cross-recognized protein is expressed in healthy tissues. Conversely, such binding could also contribute to co-targeting of additional cancer-expressed proteins if they contain a sufficiently similar epitope. Although non-specific binding has not been directly linked to adverse events in clinical studies with CAR T cells, it has been observed with TCR-engineered T cells^116^ and for therapeutic antibodies^62^.

This risk can be assessed if the epitope sequence or structure is known. For instance, a simple sequence search using the primary epitope sequence, or a structural search using Foldseek, can be applied to identify regions in other proteins that resemble the target epitope. Foldseek enables fast protein-structure searches and can therefore be used to identify proteins with local structural similarity to a known epitope^117^. Such analyses do not demonstrate binding, but they can highlight candidate off-target proteins for further evaluation.

As an example, we searched for structural similarity to the trastuzumab and pertuzumab epitopes. Both epitopes had structurally similar regions in EGFR, HER3, and HER4, which is not surprising as these receptors all belong to the ErbB family^118^. Structural alignment using PyMOL revealed close alignment of the epitope regions (**Figure 5A**), while multiple sequence alignment (MSA) showed similarity but not complete identity in the corresponding residues (**Figure 5B**). These findings indicate that related ErbB-family proteins can be flagged by structural homology searches and should therefore be evaluated as potential off-target candidates. For example, EGFR is a major target in lung cancer treatment but has been associated with various toxicities^119^.

**Figure 5.**
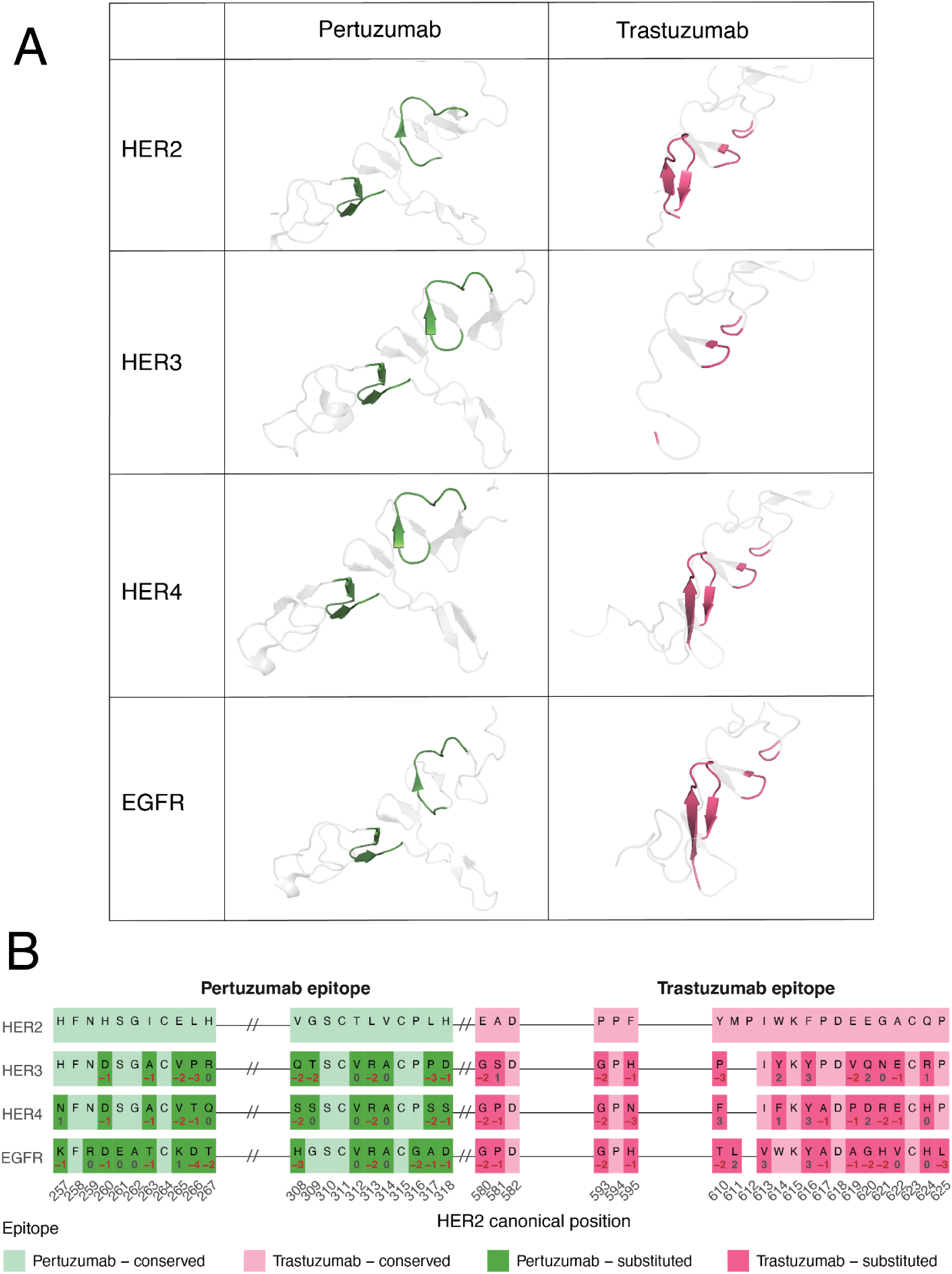
A) Structural alignment of pertuzumab and trastuzumab epitopes identified with Foldseek across HER2, HER3, HER4, and EGFR, using PyMOL. **B)** Multiple sequence alignment (MSA) of HER2, HER3, HER4, and EGFR (aligned using MAFFT), highlighting the discontinuous pertuzumab and trastuzumab epitopes. The numbers below each residue are BLOSUM62 substitution scores. Negative scores indicate less conservative amino-acid substitutions.

To assess whether this structural similarity could plausibly translate into off-target recognition, we performed an orthogonal assessment of the Foldseek hits using sequence substitution scores, backbone alignment, crystallographic flexibility, and electrostatic surface similarity (**Supplementary Table 1**). For the pertuzumab epitope, all three ErbB family members showed unfavourable amino-acid substitutions and low electrostatic similarity to HER2, despite broadly conserved backbone geometry. These results argue against substantial off-target binding risk by pertuzumab-derived CARs. For the trastuzumab epitope, the computational evidence was less uniform. EGFR showed the strongest similarity to HER2 across backbone geometry, flexibility, and electrostatic profile, whereas HER3 was partly unresolved in the available structure and therefore difficult to interpret. However, experimental studies have shown that trastuzumab does not bind EGFR, HER3, or HER4^120,121^, indicating that the cumulative sequence differences across the epitope are sufficient to prevent binding.

Together, this example illustrates that structural homology searches can identify proteins warranting further evaluation, but cannot by themselves determine cross-reactivity. Given the high sensitivity of CAR T cells, candidate off-target interactions flagged computationally should therefore be assessed experimentally during preclinical development.

## Discussion

CAR T-cell therapy has demonstrated remarkable potency due to its high sensitivity and robust cytotoxic response, and these strengths underscore the critical need for tumor-specific target selection. On-target, off-tumor toxicities can cause severe side effects, as seen with HER2-targeting CARs, where low-level HER2 expression in healthy tissues led to fatal cardiopulmonary failure^54^. Antigen selection is further complicated by the strong selective pressure exerted on targeted isoforms, which can result in isoform switch-mediated therapy-induced target loss and subsequent relapse^12^. For instance, CD19-targeting CARs in B-cell acute lymphoblastic leukemia^27^ and EGFRvIII-targeting CARs in glioblastoma multiforme^26^ have shown diminished efficacy due to the emergence of target-negative tumor clones mediated by isoform switching. These challenges highlight the importance of selecting stable, tumor-specific antigens while accounting for isoform-specific differences in subcellular localization, membrane topology, secretion, and steric accessibility of the epitope. Given the complexity of target selection, bioinformatics tools for assessing, prescreening, and identifying novel targets are needed.

Several bioinformatics pipelines for CAR target discovery have been published in recent years^25,122–124^, but current approaches remain incomplete. Gene expression-based pipelines provide broad descriptions of canonical antigen expression patterns across tissues, but generally do not resolve isoform-specific expression, which is important for assessing targetability. Gene-level approaches are also unlikely to identify completely tumor-specific targets. More recently, isoform-level expression analyses have been applied to identify tumor-specific antigens^125^. When applied to CAR target discovery, such strategies have identified the extra domain A splice variant of fibronectin as a candidate target highly expressed in tumor stromal cells, with potential to direct CAR cells into the tumor microenvironment ^126^. Despite this progress, existing pipelines generally do not assess isoform-specific protein features such as subcellular localization, membrane topology, epitope accessibility, secretion, and the potential for off-target binding. Here, we present an AI-supported framework for isoform-resolved screening of targets for CAR cell therapy or related strategies. The framework is intended to guide preclinical research and help de-risk clinical translation, but several limitations should be acknowledged. Computational target evaluation should also be interpreted in the context of increasingly flexible CAR engineering strategies. Some target liabilities may be mitigated experimentally, for example through affinity tuning, logic-gated CARs, inhibitory CARs, or engineering of the effector cell product itself. For this reason, the purpose of the workflow is not to discard all imperfect targets, but to identify molecular features that may require additional consideration during preclinical development.

First, the available data foundation is useful, but not yet optimal for CAR target evaluation. The ideal resource would be a large collection of matched tumor and healthy tissue samples profiled at the single-cell, isoform, and protein level. In practice, such datasets are not yet broadly available. Single-cell proteomics is particularly attractive because it would provide the most direct measure of target availability at the cell surface, but current methods remain insufficiently mature for large-scale and isoform-resolved applications^127,128^. Single-cell RNA sequencing using long-read or other high-sensitivity protocols offers a useful alternative by capturing transcript-level heterogeneity^129,130^, although transcript abundance is only an indirect proxy for protein abundance because of differences in translational regulation and molecule half-life^131^. This limitation is particularly relevant for CAR therapy, where the functional quantity sensed by the engineered cell is surface antigen density rather than transcript abundance. Bulk RNA sequencing, in contrast, is robust, high-throughput, and well represented in the public domain, and enables quantification of isoform fractions across tissues^132,133^, but lacks the single-cell resolution needed to resolve subclonal expression patterns. Gene-level single-cell RNA sequencing complements this by capturing cellular heterogeneity, but without transcript-level quantification it cannot distinguish between targetable and untargetable isoforms. Taken together, publicly available data remain insufficient for comprehensive CAR target assessment, but they provide a useful foundation for systematic pre-screening.

Second, many of the proposed annotations used to evaluate target suitability are based on computational predictions. Machine learning methods can estimate relevant antigen features, including subcellular localization, secretion, structural properties, antibody binding, and potential off-target interactions. These tools are useful for pre-screening, but their performance varies across proteins, prediction tasks, and biological contexts. This creates a risk of false positive predictions, which can usually be followed up experimentally, and false negative predictions, which are more difficult to identify because many features lack scalable validation methods. Thus, while computational prediction can support CAR target evaluation, it should be interpreted as a prioritization strategy rather than a substitute for experimental validation.

Third, several relevant features cannot yet be predicted with useful accuracy. Structural predictors often do not account for the effects of genetic variants, such as SNPs, on antigen stability and epitope availability. Such variants may alter the epitope sequence directly or change its structural presentation indirectly, potentially reducing CAR binding or enabling escape. Prediction of PTMs remains similarly challenging. While phosphorylation and glycosylation motifs can be predicted with moderate accuracy, the enzymatic cascades that regulate PTMs make it difficult to determine whether a given modification occurs in a specific cell type or disease context. PTMs may nevertheless affect antibody binding if located in or around the targeted epitope, and should therefore be considered when experimentally evaluating candidate targets.

Finally, the computational tools available for protein structure prediction, antibody design, affinity estimation, and protein–protein interaction modelling are advancing rapidly. As a result, specific tools highlighted here may soon be replaced by more accurate or more specialized methods. However, most of these methods are developed to optimize molecular interactions once an antigen or epitope has already been selected. They do not, by themselves, determine whether the antigen is sufficiently tumor-specific, whether the targeted epitope is retained across isoforms, whether secreted variants may act as decoys, or whether subclonal antigen-negative populations may drive relapse. The central purpose of the workflow is therefore not to prescribe a fixed set of tools, but to define the biological questions that should be considered before antibody engineering and CAR design are initiated.

Taken together, the HER2 case study illustrates how a clinically explored antigen can have target-relevant features that are not captured by gene-level expression alone. Isoform-resolved analysis identified membrane-bound and secreted variants, differences in epitope retention, subclonal expression patterns, and structurally similar regions in related proteins. Although these findings do not by themselves determine whether HER2 is suitable or unsuitable as a CAR target, they illustrate how the workflow can identify specific liabilities that should be experimentally evaluated during preclinical investigation.

Future work could extend these analyses from evaluation of individual antigens to broader screening for tumor-associated epitopes, particularly as antibody engineering expands the range of epitopes that can be targeted. The same features considered here may also be used to improve specificity by identifying molecular differences between malignant and healthy cells, for example epitopes that are retained only in cancer-associated isoforms. In addition to expression and epitope availability, the biological role of the target should be considered, as antigens that support tumor survival or proliferation may be less readily lost under therapeutic pressure. Non-canonical mechanisms of membrane association may also expand the range of candidate targets. For example, glypican-3 (GPC3), which is attached to the cell surface through a glycosylphosphatidylinositol anchor, has shown promise as a CAR target in hepatocellular carcinoma. Improved prediction and annotation of such modifications could therefore help identify antigens that are not conventional transmembrane proteins but are nevertheless accessible at the cell surface. In parallel, computational target evaluation could be combined with patient stratification, analogous to biomarker-driven treatment selection in breast cancer, to identify epitopes suitable for defined patient subsets rather than for all patients within a diagnosis. Such developments would support more tailored preclinical evaluation of CAR targets and epitopes.

## Conclusion

CAR target selection is entering a new phase. As engineered antibodies, nanobodies, de novo designed protein binders, peptide-or ligand-based binders, and other synthetic recognition domains expand the range of epitopes that can be targeted, the central limitation is shifting from whether a binding domain can be generated to whether the chosen antigen and epitope are biologically appropriate. In this study, we outline molecular features that should be considered when evaluating candidate targets for CAR cell therapy and present an in silico framework for their systematic assessment. By integrating isoform-resolved expression, subcellular localization, secretion, membrane topology, epitope mapping, structural modelling, single-cell expression analysis, and off-target assessment, the workflow extends beyond conventional gene-level target-expression screens. Using HER2 as an example, we show that clinically explored targets can have isoform-specific, soluble-decoy, subclonal, and structural features that are relevant for preclinical evaluation. Systematic evaluation of these features may help guide target and epitope prioritization as CAR design moves toward increasingly flexible and programmable antigen recognition.

## Materials and methods

### Clinical trial search

Clinical trial counts were obtained from ClinicalTrials.gov in June 2026. To estimate the overall number of CAR cell therapy trials in cancer, records were queried for interventional studies containing CAR-related terms and cancer-related indications. To estimate the number of HER2-directed CAR cell therapy trials, records were further filtered for HER2-targeting interventions and cancer indications. Trials were manually reviewed to exclude studies that did not involve CAR cell therapy or that investigated non-cancer indications. Trial status was not used as an exclusion criterion, and both completed and ongoing studies were retained.

### Isoform retrieval and filtering

Protein-coding transcript isoforms of ERBB2/HER2 were retrieved from GENCODE release 23 (GRCh38)^134^, matching the annotation used to quantify the expression data. Twelve protein-coding isoforms were identified, all of which were present in the expression matrix. Transcript identifiers, protein sequences, and annotation information were recorded. The canonical transcript, ENST00000269571, was used as the reference for epitope coordinates and isoform alignments.

### Subcellular localization prediction

Subcellular localization of the 12 protein-coding HER2 isoforms was predicted using the DeepLoc 2.1 web server^135^, applying the high-quality ProtT5-XL-UniRef50 model with extended output. Each isoform was assigned to the compartment with the highest predicted probability. Isoforms assigned to the cell membrane were retained as potentially targetable antigens, whereas secreted isoforms were retained as potential soluble decoys or antigen sinks. Isoforms assigned exclusively to intracellular compartments would have been excluded from subsequent targetability analyses; no isoform met this criterion. DeepLoc probability scores across all predicted compartments were retained for visualization and interpretation.

### Bulk isoform expression analysis

Isoform-level expression data were obtained from the UCSC Xena browser using the TcgaTargetGtex RSEM isoform TPM dataset (version 2016-09-02)^136,137^, which integrates RNA-sequencing data from TCGA, TARGET, and GTEx uniformly reprocessed against GENCODE v23. Expression of the 12 protein-coding HER2 isoforms was assessed across healthy tissues from GTEx and tumor samples from TCGA glioblastoma multiforme (n = 166), breast invasive carcinoma (n = 1,099), and stomach adenocarcinoma (n = 414). Values are distributed as log2(TPM + 0.001); these were back-transformed to TPM, negative values set to zero, and re-expressed as log2(TPM + 1) for visualization and comparison across malignant and non-malignant tissues.

### Single-cell isoform expression analysis

To assess subclonal co-expression of HER2 isoforms, three SMART-seq2 single-cell RNA-seq datasets were analyzed: GSE158631 (stomach adenocarcinoma), GSE84465 (glioblastoma), and GSE75688 (breast cancer). Raw sequencing data were retrieved from the NCBI Sequence Read Archive (SRA) using the SRA Toolkit, and transcript-level abundance was quantified using kallisto (v0.46.0) with a transcriptome index built from GRCh38.p3 (GENCODE release 23).

Cells were filtered based on library complexity (≥ 200 detected genes), mitochondrial content (≤ 15%), and sequencing depth (excluding the bottom 5th percentile), and further restricted to malignant tumor cells from primary tumor samples using dataset-specific annotations: author-assigned neoplastic cells (glioblastoma) and inferred-CNV tumor cells (breast cancer). Stomach adenocarcinoma lacked per-cell malignancy annotation, but cells were restricted to primary tumor tissue. This yielded 913 glioblastoma cells (4 patients), 60 stomach adenocarcinoma cells (3 patients) and 171 breast cancer cells (10 patients).

To evaluate therapeutic targetability, protein sequences of HER2 isoforms were extracted from the reference genome, processed through DeepLoc 2.1 for subcellular localization prediction, and mapped against known trastuzumab and pertuzumab epitopes. Isoforms were classified as “Membrane epi +” (surface-localized with a valid epitope), “Secreted epi +” (extracellular with a valid epitope), or “Non-targetable” (lacking valid epitopes or surface localization). Transcript counts were binarized (detected = nonzero estimated count), and per-sample expression profiles across these categories were visualized using pheatmap v1.0.13 ^138^.

### Membrane topology prediction

Transmembrane topology of the retained protein-coding HER2 isoforms was predicted using the DeepTMHMM 1.0 web server^101^ with default settings. Predicted transmembrane helices and extracellular, intracellular, and signal-peptide regions were used to determine whether epitope-containing sequences were located in extracellular regions accessible to CAR binding. Isoforms lacking a predicted transmembrane helix were evaluated as potential soluble variants, whereas isoforms containing a transmembrane helix were assessed for the size and orientation of their predicted extracellular domains.

### Epitope identification and mapping

Known trastuzumab and pertuzumab epitopes were identified through a literature search assisted by the large language models Claude (Anthropic) and Gemini (Google). The models were used to locate candidate publications and reported epitope coordinates; all residue positions and source references were subsequently verified manually against the original publications. Verified epitope residues were mapped to the canonical HER2 protein sequence, ENST00000269571. The HER2 isoforms were aligned using MAFFT v7^139^ (with arguments --localpair --maxiterate 1000), and the corresponding epitope positions were transferred through the alignment to assess epitope retention across isoforms.

### Three-dimensional structure prediction

Three-dimensional structures of the retained protein-coding HER2 isoforms were predicted using the AlphaFold 3 web server ^106^ in template-free mode. Five models were generated for each isoform. As the predicted structures were consistent across seeds, the top-ranked model was selected for downstream analysis. Structures were visualized in PyMOL v3.1.6.1 ^140^, and the mapped trastuzumab and pertuzumab epitope residues were inspected to assess their spatial presentation and potential steric accessibility.

### Off-target binding assessment

Structural similarity between the HER2 epitopes and regions in other human proteins was assessed using Foldseek (release v10-941cd33, default parameters) to identify potential off-target candidates. As Foldseek indexes canonical protein sequences, this analysis was performed at the gene level rather than the isoform level. Canonical protein sequences for HER2 (ENST00000269571), HER3 (ENST00000267101), HER4 (ENST00000342788), and EGFR (ENST00000275493) were aligned using MAFFT v7 with `--localpair --maxiterate 1000`. Amino-acid substitutions at the pertuzumab and trastuzumab epitope positions were scored using the BLOSUM62 substitution matrix, with negative scores indicating less conservative substitutions.

Crystal structures of HER2 in complex with pertuzumab (PDB: 1S78) and trastuzumab (PDB: 1N8Z), together with unbound structures of HER3 (PDB: 1M6B), HER4 (PDB: 2AHX), and EGFR (PDB: 1NQL), were retrieved from the Protein Data Bank. Solvent molecules and small-molecule ligands were removed, and duplicate chains in the asymmetric unit were excluded. For each antibody epitope, the corresponding regions in HER3, HER4, and EGFR were aligned to HER2 in PyMOL v3.1.6.1. Backbone RMSD was calculated from Cα atoms without outlier rejection (`cycles = 0`) to quantify local structural similarity across the aligned epitope regions.

For the pertuzumab epitope, structural alignment was performed using HER2 residues 235–245 and 286–296 in 1S78 chain A, corresponding to canonical HER2 positions 257–267 and 308–318. The equivalent regions were HER3 residues 229–239 and 279–289, HER4 residues 226–236 and 276–286, and EGFR residues 229–239 and 280–290. For the trastuzumab epitope, HER2 residues 558–560, 571–573, and 588–603 in 1N8Z chain C, corresponding to canonical positions 580–582, 593–595, and 610–625, were aligned to HER3 residues 550–552, 563–565, and 580–593; HER4 residues 548–550, 561–563, and 578–591; and EGFR residues 551–553, 564–566, and 581–595.

Crystallographic B-factors were extracted for resolved epitope Cα atoms and used as an approximate measure of local conformational mobility. Residues with B-factor values of 0.0 were treated as unresolved and excluded. B-factor comparisons were restricted to the unbound HER3, HER4, and EGFR structures, which were of similar resolution, while the antibody-bound HER2 structures were excluded from cross-protein B-factor comparisons because complex formation may influence local mobility.

Electrostatic potentials were calculated using PDB2PQR v3.7.1^141^ with the AMBER force field and APBS v3.4.1. Potentials were sampled at epitope Cα positions by trilinear interpolation of the APBS potential grid. Equivalent positions across ERBB-family members were matched through the MAFFT alignment, with crystallographic residue numbering reconciled to the corresponding canonical sequence positions. Electrostatic similarity was quantified as the Pearson correlation between per-residue potential profiles for HER2 and each family member across positions resolved in both structures. This analysis provides a coarse comparison of the electrostatic environment of the epitope rather than a direct estimate of antibody-binding affinity or surface complementarity.

## Data availability

All datasets used in this study are publicly available. Bulk transcript-level expression data were obtained from the UCSC Xena TCGA/GTEx resource, and single-cell RNA-sequencing datasets were retrieved from the NCBI Sequence Read Archive under the accessions described above. Protein structures were obtained from the Protein Data Bank. Derived data used to reproduce the HER2 analyses are available at: https://github.com/biosurf/biosurf/tree/master/cartcontent.

## Code availability

Code used for the analyses presented in this study is available at: https://github.com/biosurf/biosurf/tree/master/cartcontent. The repository also contains vignettes describing the major analysis steps and can be adapted to evaluate additional CAR targets and updated reference datasets.

## Funding

LRO was funded by the Independent Research Fund Denmark (grant 8048-00078B) and the Novo Nordisk Foundation (grant NNF22OC0071681). The LEO Foundation Center for Skin Immunology Research is supported by LEO Fondet (grant LF18500). NIVI Research Center is part of The Novo Nordisk Foundation Initiative on Vaccines and Immunity (NIVI) and is supported by the Novo Nordisk Foundation (grant NNF23SA0088562). JHdC received funding from the European Union’s Horizon Europe programme (grant 101169223). MBB was funded by the Lundbeck Foundation (grant R381-2021-1278 and R480-2024-1167). KVS was funded by Lundbeck Foundation (grant R413-2022-878).

## Author contributions

LRO and MBB conceptualized the study. GM, LVJ, JHdC, MHH, and AD performed the data analysis. All authors interpreted the results. LRO, MBB, and GM wrote the manuscript. All authors reviewed and approved the manuscript.

## Competing interests

MBB has received consulting honorariums from Janssen and Kite/Gilead.

## Supplementary Material

**Supplementary Figure 1.**
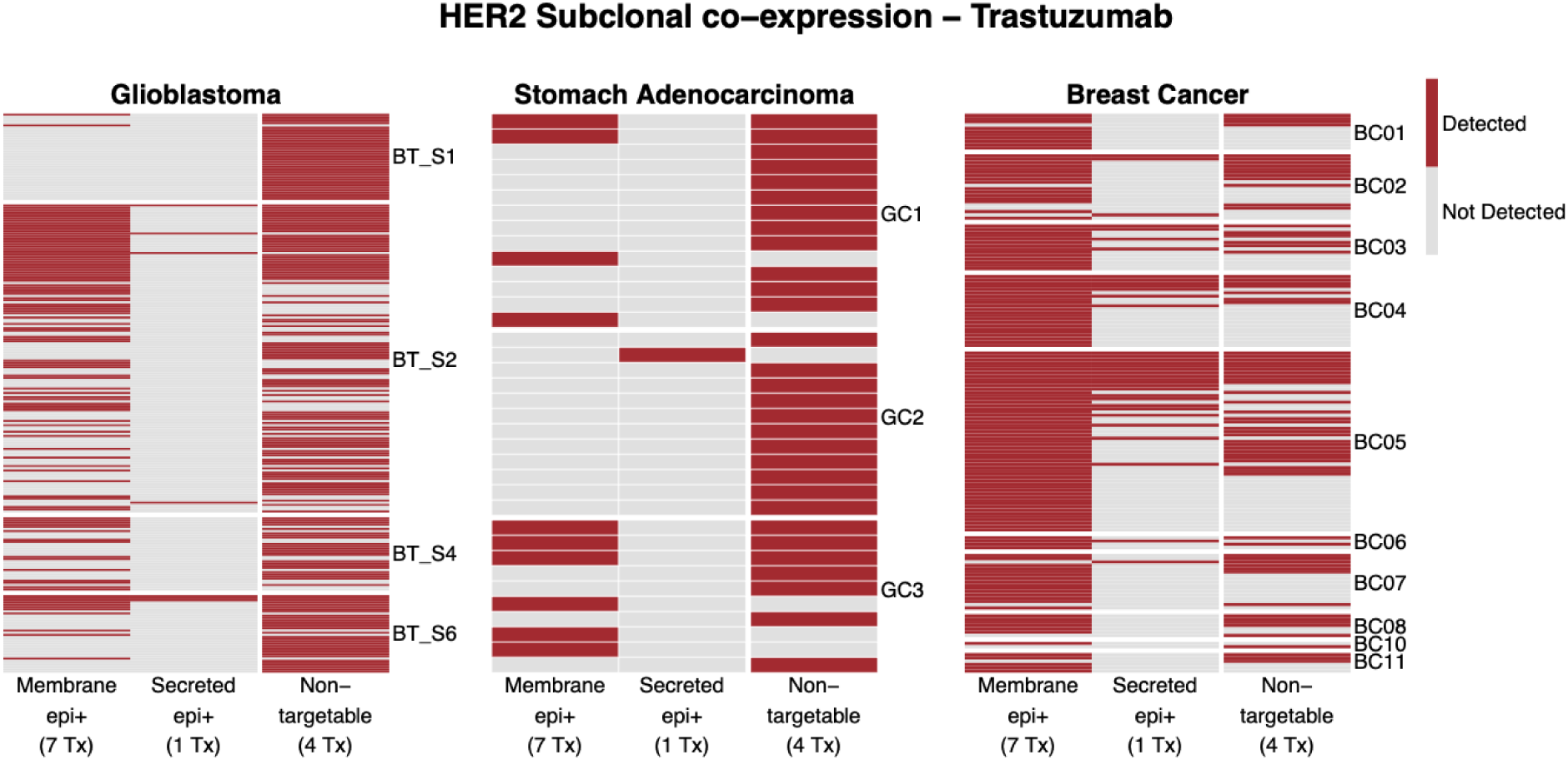
Single-cell expression of HER2 isoforms classified by trastuzumab targetability. Binary heatmap showing detection of HER2 transcript categories across single cells from glioblastoma, stomach adenocarcinoma, and breast cancer. Each row represents an individual cell, grouped by patient; columns indicate detection or absence of at least one transcript classified as membrane-bound epitope-positive, secreted epitope-positive, or non-targetable for trastuzumab. Only cells with detectable HER2 transcripts are shown. Cells without detectable HER2 expression were excluded from visualization: GBM, n = 660; stomach adenocarcinoma, n = 24; breast cancer, n = 13. The number of transcripts assigned to each category is indicated in parentheses. The same underlying filtered single-cell datasets were used as for Figure 4B.

---

**Supplementary Table 1.**
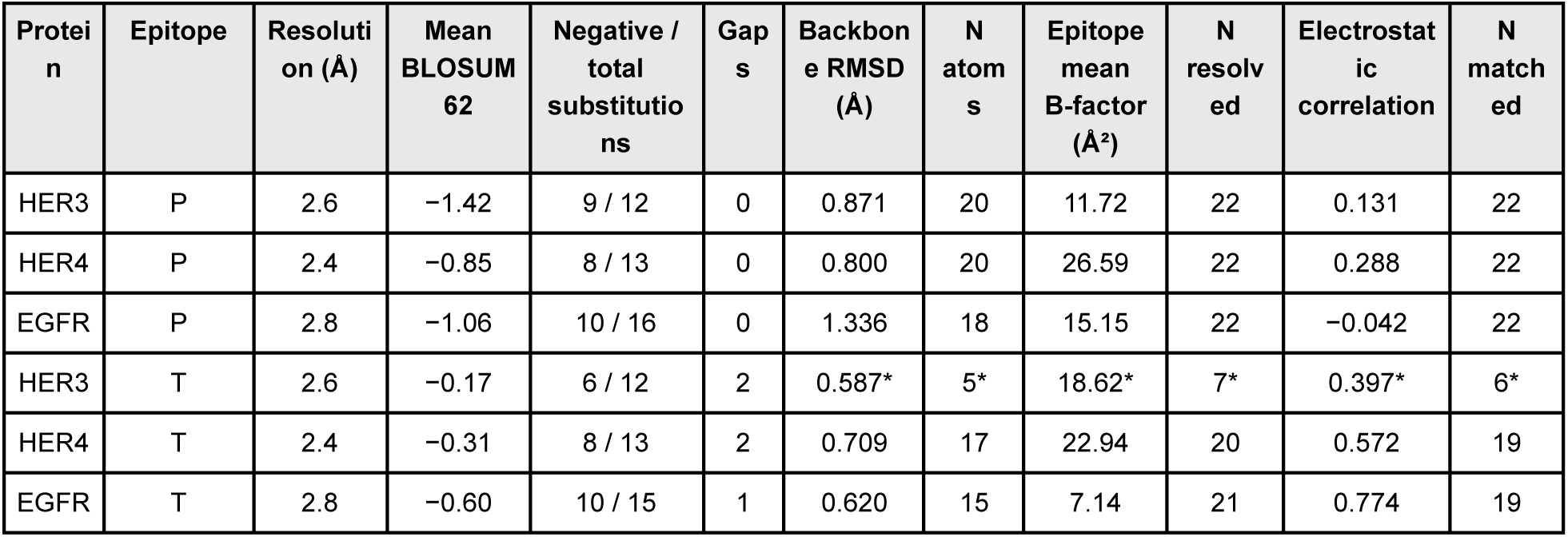
Multi-parameter assessment of structurally similar ERBB-family epitopes. Foldseek-identified regions in HER3, HER4, and EGFR were compared with the corresponding HER2 epitopes recognized by pertuzumab and trastuzumab. HER2 served as the reference and is therefore not shown. Each epitope comprises 22 canonical HER2 positions. Resolution refers to the structure used for B-factor analysis; absolute B-factor values were compared only across the three unbound structures, which were of similar resolution (2.4–2.8 Å). Mean BLOSUM62 scores were calculated across substituted positions only, excluding identical residues and gaps. Negative/total substitutions indicates the number of substitutions with negative BLOSUM62 scores out of all substituted positions. Gaps indicate canonical HER2 epitope positions absent from the corresponding ERBB-family sequence in the MAFFT alignment. Backbone RMSD was calculated from aligned Cα atoms without outlier rejection (cycles = 0) after structural alignment of the epitope regions in PyMOL; N atoms indicates the number of Cα pairs included. Epitope mean B-factor was calculated across resolved epitope Cα atoms, excluding positions with B = 0.0; N resolved indicates the number of residues included. Electrostatic correlation is the Pearson correlation between APBS-derived per-residue electrostatic potentials at matched epitope Cα positions in HER2 and the corresponding ERBB-family member; N matched indicates the number of aligned positions resolved in both structures. The correlation reflects similarity in the pattern of electrostatic potential rather than its absolute magnitude, and Cα sampling provides a coarse measure of the local backbone electrostatic environment rather than side-chain contact chemistry. Asterisks denote values based on too few resolved positions for reliable interpretation; thirteen consecutive residues are unresolved in HER3 at the trastuzumab epitope.

## Notes

https://biosurf.org/cart.html

https://transcript-explorer-app.onrender.com/

