## Supplemental figure 1 for "AI supported *in silico* screening of chimeric antigen receptor therapy targets"

### Supplementary Material

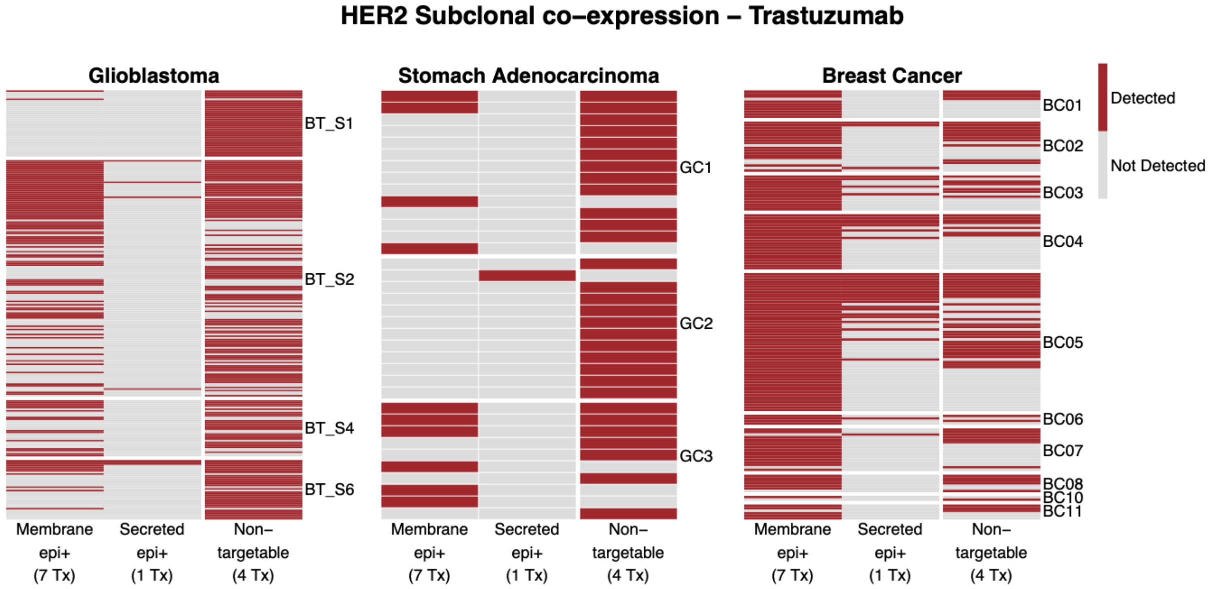


**Supplementary Figure 1**. Single-cell expression of HER2 isoforms classified by trastuzumab targetability. Binary heatmap showing detection of HER2 transcript categories across single cells from glioblastoma, stomach adenocarcinoma, and breast cancer. Each row represents an individual cell, grouped by patient; columns indicate detection or absence of at least one transcript classified as membrane-bound epitope-positive, secreted epitope-positive, or non-targetable for trastuzumab. Only cells with detectable HER2 transcripts are shown. Cells without detectable HER2 expression were excluded from visualization: GBM, n = 660; stomach adenocarcinoma, n = 24; breast cancer, n = 13. The number of transcripts assigned to each category is indicated in parentheses. The same underlying filtered single-cell datasets were used as for **Figure 4B**.
