## Supplemental table 1 for "AI supported *in silico* screening of chimeric antigen receptor therapy targets"

**Supplementary Table 1**. Multi-parameter assessment of structurally similar ERBB-family epitopes. Foldseek-identified regions in HER3, HER4, and EGFR were compared with the corresponding HER2 epitopes recognized by pertuzumab and trastuzumab. HER2 served as the reference and is therefore not shown. Each epitope comprises 22 canonical HER2 positions. Resolution refers to the structure used for B-factor analysis; absolute B-factor values were compared only across the three unbound structures, which were of similar resolution (2.4–2.8 Å). Mean BLOSUM62 scores were calculated across substituted positions only, excluding identical residues and gaps. Negative/total substitutions indicates the number of substitutions with negative BLOSUM62 scores out of all substituted positions. Gaps indicate canonical HER2 epitope positions absent from the corresponding ERBB-family sequence in the MAFFT alignment. Backbone RMSD was calculated from aligned Cα atoms without outlier rejection (cycles = 0) after structural alignment of the epitope regions in PyMOL; N atoms indicates the number of Cα pairs included. Epitope mean B-factor was calculated across resolved epitope Cα atoms, excluding positions with B = 0.0; N resolved indicates the number of residues included. Electrostatic correlation is the Pearson correlation between APBS-derived per-residue electrostatic potentials at matched epitope Cα positions in HER2 and the corresponding ERBB-family member; N matched indicates the number of aligned positions resolved in both structures. The correlation reflects similarity in the pattern of electrostatic potential rather than its absolute magnitude, and Cα sampling provides a coarse measure of the local backbone electrostatic environment rather than side-chain contact chemistry. Asterisks denote values based on too few resolved positions for reliable interpretation; thirteen consecutive residues are unresolved in HER3 at the trastuzumab epitope.

| Protein | Epitope | Resolution (Å) | Mean BLOSUM62 | Negative / total substitutions | Gaps | Backbone RMSD (Å) | N atoms | Epitope mean B-factor (Å²) | N resolved | Electrostatic correlation | N matched |
|---|---|---|---|---|---|---|---|---|---|---|---|
| HER3 | P | 2.6 | −1.42 | 9 / 12 | 0 | 0.871 | 20 | 11.72 | 22 | 0.131 | 22 |
| HER4 | P | 2.4 | −0.85 | 8 / 13 | 0 | 0.800 | 20 | 26.59 | 22 | 0.288 | 22 |
| EGFR | P | 2.8 | −1.06 | 10 / 16 | 0 | 1.336 | 18 | 15.15 | 22 | −0.042 | 22 |
| HER3 | T | 2.6 | −0.17 | 6 / 12 | 2 | 0.587* | 5* | 18.62* | 7* | 0.397* | 6* |
| HER4 | T | 2.4 | −0.31 | 8 / 13 | 2 | 0.709 | 17 | 22.94 | 20 | 0.572 | 19 |
| EGFR | T | 2.8 | −0.60 | 10 / 15 | 1 | 0.620 | 15 | 7.14 | 21 | 0.774 | 19 |
